# A Lenalidomide-Inducible Suicide Switch for Gene- and Cell-Based Therapy

**DOI:** 10.64898/2026.09.30.755642

**Authors:** Ditsa Sarkar, William Lin, Nelson H. Knudsen, Jalen Evans, Joanna Kim, Stephen M. Coscia, Isabel C. Lane, Andrew Cheng, Noor Radde, Manuella Talla, Angelique Pham, Tamina Kienka, Emily Schneider, Gabriele Kembuan, Mikołaj Słabicki, Vidyasagar Koduri, Adam S. Sperling, Marcela V. Maus, Robert T. Manguso, Max Jan

## Abstract

Genetically modified immune effector cells, including chimeric antigen receptor (CAR) T cells, have emerged as transformative anti-cancer agents. However, these autonomous, living therapies are limited by difficult-to-control hyperactivation toxicities. To achieve their full potential as therapies for a broader range of diseases, CAR T cells must target novel antigens and integrate further potency enhancements that raise the possibility of novel and intensified toxicities. High performance safety systems are needed to restrain these highly active next-generation gene- and cell-based therapies. Cell therapy suicide switches remain infrequently used, and are currently limited by inaccessible controller drugs, incomplete cell depletion, fitness cost, and immunogenicity. To address these issues, we engineered a fully human sequence-based cell therapy suicide switch based on the overexpression of Caspase-Activated DNase (CAD) and its stoichiometric chaperone/inhibitor ICAD fused to a lenalidomide-inducible degron. Acute destabilization of ICAD with the broadly available, off-patent molecular glue degrader drug lenalidomide uncaged CAD to induce DNA damage and cell death. By this degradable cage mechanism, the ICAD-CAD suicide switch achieved long-term expression in primary human T cells without detectable leakiness, and deep drug-induced cell depletion. The ICAD-CAD switch was generalizable to alternative acute destabilization systems. Genome-wide CRISPR screening established the necessity of CAD and the lenalidomide-interacting E3 ubiquitin ligase substrate adaptor CRBN for switch function. When co-delivered with a CAR, ICAD-CAD CAR T cells retained effector functions and could be rapidly depleted with lenalidomide.

## INTRODUCTION

CAR T cell therapies are now established medical breakthroughs in the treatment of multiple B cell cancers and have shown promising activity signals in early-stage clinical trials for select solid tumors, T cell malignancies, and autoimmune diseases^1,2^. Yet the therapeutic window of these living drugs remains limited. While common class-wide complications of CAR T cell therapy including cytokine release syndrome and immune effector cell-associated neurotoxicity syndrome are manageable in the vast majority of cases, rare toxicities including immune effector cell-associated hemophagocytic lymphohistiocytosis-like syndrome and colitis can be difficult to manage^3,4^. As CAR T cell therapies are advanced in new disease applications, severe ON-target OFF-tumor toxicities can arise unexpectedly in early stage clinical trials^5^. Additional genetic modifications can dramatically enhance CAR T cell potency and may be needed for solid tumor targeting^6–8^, however these functional enhancements may also lead to new or exacerbated toxicities. In high-risk scenarios, definitive safety systems are needed to intercept severe complications of CAR T cell therapy.

Suicide switches for cell therapy were first developed over three decades ago, yet their clinical use remains limited. Early designs such as the herpes simplex virus thymidine kinase/ganciclovir system employed non-human proteins, and were susceptible to T cell recognition of the foreign sequences and immune rejection^9^. The most commonly used systems are 1) the expression of surface epitopes that can be targeted with approved antibody-based therapeutics^10,11^ and 2) inducible Caspase 9 (iCasp9), wherein a dimerizer small molecule AP1903 activated engineered iCasp9 proteins^12^. However, the expression of monoclonal antibody epitopes relies on non-cell-intrinsic death mechanisms that can be dysfunctional in patients with advanced cancers, leading to incomplete cell removal. Chemically-inducible dimerization systems can be susceptible to narrow expression ranges for optimal function, with high expression leading to leaky drug-independent dimerization and low expression leading to insufficient drug-induced dimerization^13,14^. Indeed, although iCasp9 has been successfully used to intercept toxicities in clinical trials, iCasp9+ cells have escaped switch activation and re-expanded^12,15^. Furthermore, AP1903 is not clinically approved, thus iCasp9 is unavailable to the majority of cell therapy translational efforts. The breadth of pre-clinical safety systems has been recently reviewed^16^. There is an unmet need for more ideal suicide switch systems that 1) are composed of human sequences, 2) fit existing vector and manufacturing constraints, 3) do not reduce engineered cell fitness, and 4) completely eliminate engineered cells upon induction by a 5) broadly available drug controller. Such systems may become important safeguards for investigational cell therapies that incorporate novel designs, are highly active, and/or are intended for patient populations at elevated risk for severe toxicity.

Here we present the development of a suicide switch that works by a fundamentally distinct mechanism. Inspired by natural cell death systems composed of stoichiometric pairs of pro- and anti-apoptotic factors that are tonically expressed without leakiness, we envisioned a suicide switch composed of a multi-cistronic transgene pair of a pro-apoptotic protein restrained by an anti-apoptotic protein that could be acutely destabilized under user control. We hypothesized that, if the inhibitory protein were sufficiently effective, arbitrarily high transgene expression could be achieved without leakiness, and induced degradation of the inhibitor could uncage high concentrations of the effector for complete cell elimination. Caspase-activated DNase (CAD) is functionally sequestered by its stoichiometric inhibitor (ICAD)^17,18^. ICAD serves as a co-translational chaperone for CAD to cage newly folded CAD molecules in an inert heterodimer^19–21^. Upon apoptosis induction, Caspase 3 cleaves ICAD, releasing CAD to homodimerize, cleave genomic DNA, and induce cell death^22^. Previously, avian and murine Cad and Icad, the latter fused to an auxin-induced degron, had been overexpressed to generate a cell death switch triggered by the plant hormone and molecular glue degrader auxin^23^. This switch, although composed of disparate parts, demonstrated that acute destabilization of ICAD with a degron tag could uncage CAD and induce cell death in mammalian cells.

Targeted protein degradation has emerged as a mainstream therapeutic approach, including the development of novel classes of protein degraders and the serendipitous discovery that approved anti-cancer drugs such as lenalidomide act as molecular glues to degrade disease-relevant proteins^24^. Lenalidomide is a frontline, off-patent therapy for people with hematologic malignancies including multiple myeloma and myelodysplastic syndrome with deletion of chromosome 5q^25^. In broad clinical testing, lenalidomide has been found to be safe but with limited efficacy in people with advanced solid tumors, providing rationale for lenalidomide as an approved, broadly available, and safe CAR T cell controller drug^25^. Lenalidomide acts as a molecular glue to recruit disease-relevant neosubstrate proteins to the CRL4_CRBN_ E3 ubiquitin ligase for subsequent proteasomal degradation^26,27^. Engineered zinc finger domains from neosubstrate proteins can be used as degron tags for lenalidomide-inducible degradation of CARs and other fusion proteins^28–32^. Here, we report the development of a fully human sequence-based suicide switch based on lentiviral overexpression of CAD and zinc finger degron-tagged ICAD. This ICAD-CAD suicide switch was optimized for long-term expression in human T cells, and demonstrated best-in-class cell depletion upon lenalidomide treatment. Mechanistically, we used chemogenetic and genome-scale CRISPR screening approaches to establish that the DNase-based switch was sensitive to CAD concentration and required CRL4_CRBN_, but otherwise was insensitive to loss of DNA damage response or cell death machinery. When co-delivered with a CAR, the ICAD-CAD suicide switch could rapidly deplete CAR T cells and prevent their effector functions.

## RESULTS

### Design of an ICAD-CAD suicide switch

We envisioned a human sequence-based ICAD-CAD suicide switch wherein 1) co-overexpression of ICAD and CAD would generate inert ICAD/CAD heterodimers until 2) acute pharmacologic destabilization of ICAD would release CAD to induce cell death (Fig. 1A). To engineer a degradable variant of ICAD, we used a lenalidomide-responsive zinc finger-based degron tag that has been used to regulate the abundance of CARs, cytokines, Cas9, and anti-CRISPR proteins^28,30,32,33^ and generated a GFP-ICAD-degron fusion protein (Fig. 1B). We evaluated the dose response and kinetics of GFP-ICAD-degron depletion in Jurkat T cells and compared the specificity of the degron with other cereblon E3 modulatory drugs (CELMoDs) using DMSO control. The half-maximal concentration of lenalidomide, pomalidomide and mezigdomide for GFP-ICAD-degron depletion was ∼0.6 nM, 0.2 nM and 0.9 pM respectively (Fig. 1B), and half-maximal protein degradation occurred in less than one hour in all three cases (Fig. 1C). This suggests that the ICAD-degron fusion protein could be depleted by diverse cereblon E3 modulatory drugs.

**Figure 1.**
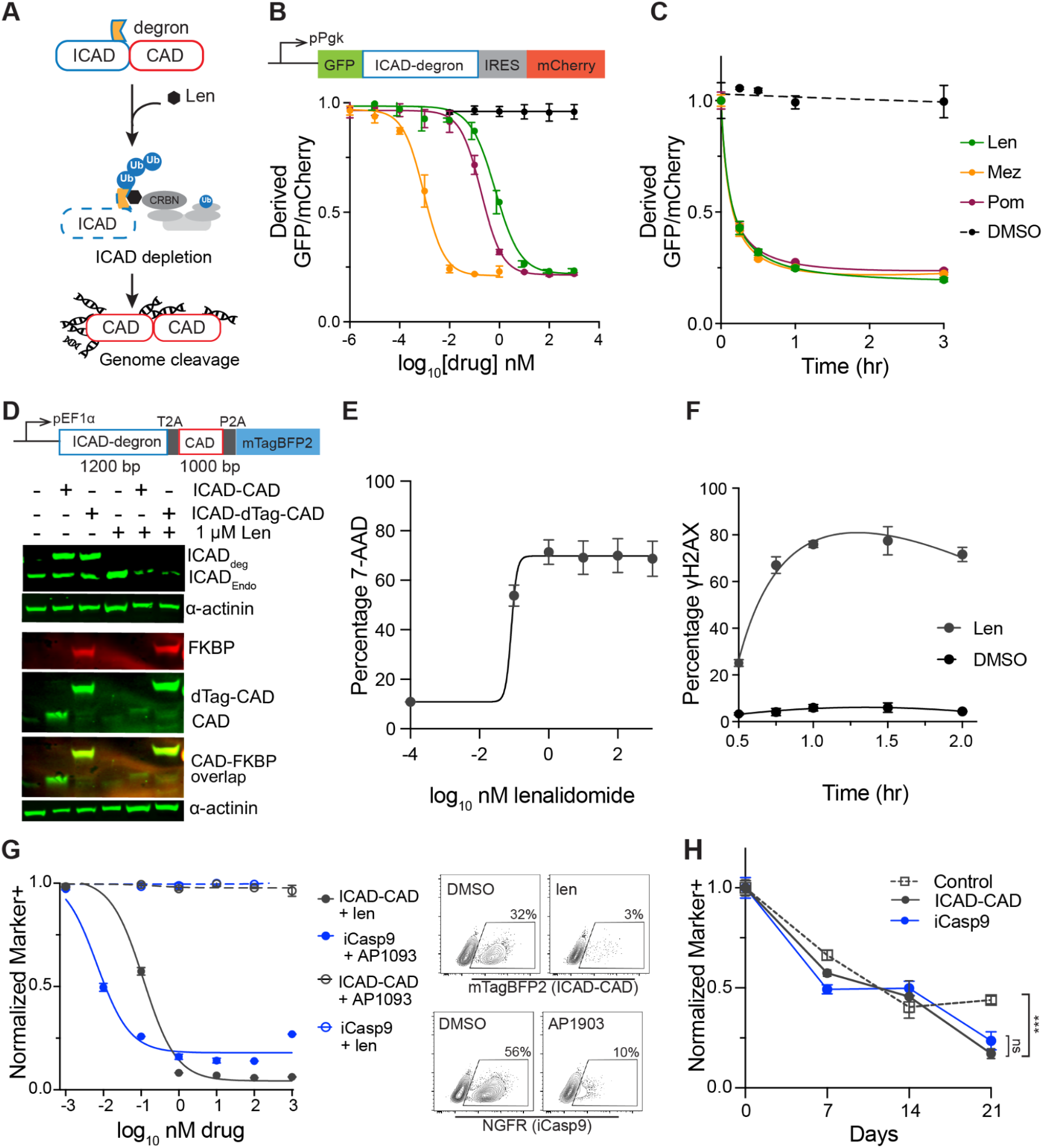
Development of a lenalidomide-inducible ICAD-degron-CAD suicide switch (A) Schematic of ICAD-CAD suicide switch mechanism. Lenalidomide (len) recruits E3 ubiquitin ligase complex to degron-tagged ICAD resulting in CAD-mediated DNA double strand breaks. (B) Vector map of a lenalidomide-inducible ICAD-degron-GFP with a fluorescent mCherry reporter. Plot showing degradation of GFP-ICAD-degron fusion protein in Jurkat T cells after treatment with varying lenalidomide (len), mezigdomide (mez) and pomalidomide (pom) concentrations. Y-axis displays a derived GFP-mCherry fluorescence ratio calculated from FlowJo software using the formula, Ratio= (GFP/mCherry). (C) Depletion of GFP-ICAD-degron fusion protein depletion after treatment with 1 µM len, pom and 1 nM mez and DMSO control in Jurkat T cells at different time points. Derived GFP-mCherry ratio was calculated as described above. (D) Construct map of a lenalidomide-inducible ICAD-degron-CAD suicide switch with a fluorescent mTagBFP2 reporter. Western blot of ICAD-CAD suicide switch, ICAD-dTag-CAD and untransduced (UTD) Jurkat T cells with or without 1 µM lenalidomide after 2 hr of treatment. anti-FKBP was used to detect dTAG^34^. (E) Percentage of 7AAD in primary human T cells expressing ICAD-CAD suicide switch after 20 hr treatment with varying lenalidomide concentrations. (F) Percentage of γ-H2AX in primary human T cells expressing ICAD-CAD suicide switch after 1 µM lenalidomide treatment at different time points. (G) Left panel shows dose response of ICAD-CAD and iCasp9 suicide switches at different drug concentration. Primary human T cells containing ICAD-CAD or iCasp9 systems were individually treated with lenalidomide or AP1903 at various concentrations for 24 hr. Marker+ population was gated from live cells as shown in Fig. S1B. Right panel shows a representative Flow plot of mTagBFP2 (ICAD-CAD suicide switch) at the top and NGFR (iCasp9 suicide switch) at the bottom. Cells were normalized to the respective untreated population. (H) Long-term proliferation assay of ICAD-CAD and iCasp9 suicide switches at given time points. Primary human T cells were transduced with either of the suicide switches and periodic flow measuremets of their marker (BFP or NGFR) were carried out and compared to a control. Cells were normalized to day 0 marker intensity. Flow gating is depicted in Fig. S1B.Panel B-C performed with technical duplicate. Error bars represented as mean ± SD. Panel E-F performed with 2 Healthy donors (HD), mean ± SD. Panel G carried out with 1HD, technical duplicate, mean ± SD. Panel H with 1HD, technical triplicate, statistical analysis by 2-way ANOVA with Šídák’s multiple comparisons test at day 21 (\**P* < 0.05, \*\**P* < 0.01, \*\*\**P* < 0.001). Donor 2 data shown Figure SF2.

Having established drug-inducible depletion of an ICAD-degron fusion protein, we then generated a putative suicide switch composed of single lentiviral construct encoding ICAD-degron, CAD and mTagBFP2 as a multicistronic gene each separated by ribosome skipping elements (Fig. 1D). To directly measure the abundance of ICAD and CAD, we performed immunoblots. To distinguish between endogenous and over-expressed CAD, we fused CAD to dTag^34^ and expressed them with ICAD-degron (Fig. S1A). We detected ICAD-degron and CAD depletion upon lenalidomide treatment (Fig. 1D). To determine the effect of ICAD and CAD overexpression on T cell survival with and without lenalidomide, we transduced primary T cells with a series of ICAD-CAD lentivectors differentiated by the CAD transgene sequence (Fig. S2A). We reasoned that the native and codon optimized CAD sequences would have different translation rates and stoichiometries of ICAD and CAD proteins. The CAD p.R11W mutation was an accidental event in lentivector cloning and was included in preliminary experiments. Upon lenalidomide administration, the majority of transduced cells were depleted with all iterations (Fig. S2A). Whereas the codon optimized CAD- and native CAD-based suicide switches reduced the relative fitness of the ICAD-CAD transduced cells, the CAD p.R11W mutant construct did not reduce cell fitness (Fig. S2A). Modeling of the CAD p.R11W substitution in the context of the ICAD-CAD heterodimer structure^21^ places the substitution in the Cell death-Inducing DFF45-like Effector (CIDE) domain near the protein-protein interface (Fig. S2B). Additional structurally adjacent substitutions of CAD p.R11W-interacting residues also rescued the fitness cost of ICAD-CAD co-expression in Jurkat T cell line (Fig. S2C), suggesting that modulation of the CAD CIDE domain adjacent to the protein-protein interaction interface can enhance the tolerability of its overexpression. We used the ICAD-CAD p.R11W suicide switch for all subsequent studies, which will be referred to as the ICAD-CAD suicide switch below.

We next evaluated the function of the ICAD-CAD suicide switch in primary T cells. Half-maximal induction of cell death, as quantified by 7AAD staining, was observed with sub-nanomolar lenalidomide concentrations (Fig. 1E) and induced cell death was predominantly apoptotic as confirmed by Annexin V/7AAD experiment (Fig. S3). The peak plasma concentration of lenalidomide in people receiving standard dosing for multiple myeloma is in the micromolar range^35^. Half-maximal DNA damage occurred within one hour of drug exposure (Fig. 1F). In summary, subnanomolar lenalidomide concentrations rapidly induced the death of T cells expressing the ICAD-CAD suicide switch.

iCasp9 is currently the most used safety switch in cell therapy clinical trials, therefore we performed a head-to-head comparison between iCasp9 and ICAD-CAD. The ICAD-CAD switch demonstrated more complete depletion of engineered cells versus the iCasp9 system across a broad rimiducid concentration range, with an upper boundary Hook effect for iCasp9 indicated by reduced efficacy in the micromolar range (Fig. 1G and Fig. S4A). In a long-term proliferation assay, we found that ICAD-CAD was comparable to or outcompeted iCasp9 in different T cell donors (Fig. 1H, Fig. S4B). Overall, utilizing all-human sequences and a clinically approved drug controller, we developed a candidate lenalidomide-inducible suicide switch that rapidly induced cell death with deeper induced cell depletion than the well-validated iCasp9 system.

### Generalizable chemogenetic ICAD-CAD switches

To evaluate the robustness and generalizability of the ICAD-CAD switch with other conditional degradation systems, we fused ICAD to dTag^34^ and SMASh tag^36^ and expressed them with CAD. In Jurkat T cells, dTag-13 induced dose-dependent depletion of ICAD-dTag-CAD-expressing cells (Fig. 2A, Fig. S1A). By contrast, the ICAD-SMASh-CAD switch, which is designed to shut off accumulation only of newly translated ICAD-SMASh tag fusion proteins, did not induce cell death with the addition of asunaprevir. These data, together with prior auxin-induced degron fusions with murine ICAD^23^, demonstrate that molecularly diverse chemogenetic systems that act by rapidly destabilizing ICAD induce cell death in the context of CAD overexpression.

**Figure 2.**
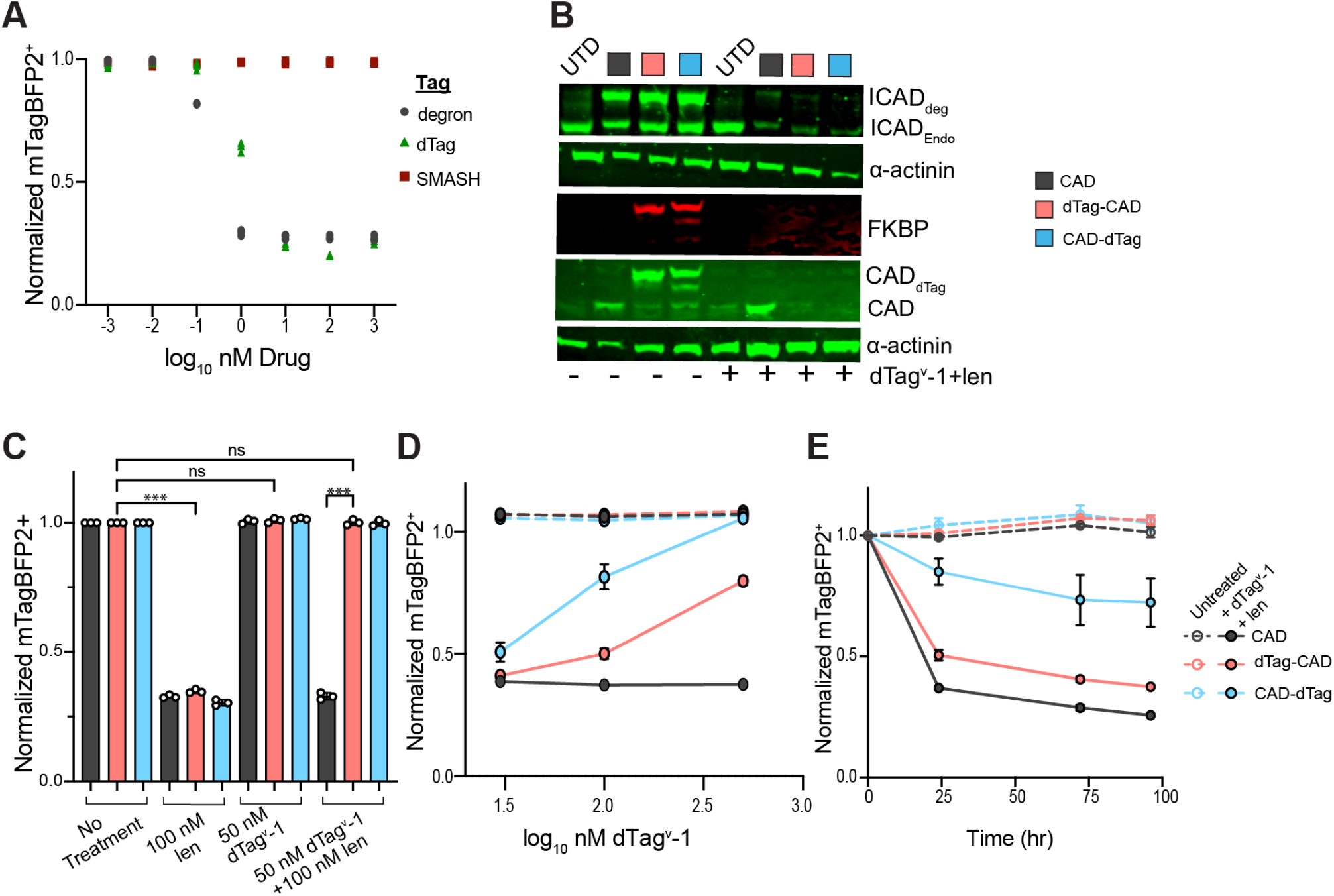
ICAD-CAD switch generalizability and sensitivity to CAD modulation mTagBFP2 intensity of ICAD-CAD switches expressed in Jurkat T cells measured by Flow cytometry. Representative Flow gating shown in Fig. S1B. Normalization of mTagBFP2 was performed with the respective untreated population. (A) ICAD fused to different conditional degradation systems and expressed with CAD. Normalized mTagBFP2 intensity of ICAD-CAD switches treated with their respective chemical inducer for 48 hours. (B) Western blot of CAD fused to dTag at N- or C-terminal and expressed with ICAD with or without 100 nM dTag^v^-1 + 500 nM len after 2 hr of treatment. (C) Normalized mTagBFP2 intensity of cells transduced with different ICAD-CAD inducible systems treated with or without 50 nM dTag^v^-1 and 100 nM Len. (D) Plot of mTagBFP2 intensity of ICAD-CAD inducible systems after 20 hr treatment with varying concentration of dTag^v^-1 and 100 nM len. (E) Depletion of mTagBFP2 expression of different ICAD-CAD systems treated with 100 nM dTag^v^-1 and 100 nM len at different time points. mTagBFP2+ normalized with untreated condition at t=0. Panels A and C-E performed in technical triplicate, mean ± SD . Panel C analyzed by ordinary one-way ANOVA with Tukey’s multiple comparisons test (\**P* < 0.05, \*\**P* < 0.01, \*\*\**P* < 0.001)

We next evaluated whether the ICAD-CAD switch was sensitive to modulation of CAD concentration. To this end, we created tricistronic constructs encoding (i) ICAD fused to the lenalidomide-responsive degron (ii) CAD fused to dTag on either its N- or C-terminus (hereafter, named dTag-CAD or CAD-dTag respectively) and (iii) mTagBFP2 (Fig. S1A). We observed that treatment with the VHL-recruiting PROTAC dTag^v^-1, induced dTag-CAD or CAD-dTag degradation (Fig. 2B). Furthermore, simultaneous co-administration of dTag^v^-1 and lenalidomide, rescued lenalidomide-induced cell death (Fig. 2C). In a dose response experiment, titration of dTag^v^-1 versus a constant lenalidomide dose tuned the amount of cell depletion (Fig. 2D). In a time course experiment, the majority of the cell depletion occurred within the first 24 hours (Fig. 2E). In summary, using dTag-CAD and dTag^v^-1, we established CAD protein abundance as a critical modulator of suicide switch performance. In the future, beyond cell therapy applications, fusions of CAD to proteins of interest may be used as stability reporters coupled to the lenalidomide-inducible cell depletion switch, in order to pan for protein degraders with a fast and sort-free viability-based screening assay^37^.

### Identification of resistance mechanisms to the ICAD-CAD suicide switch

To identify potential mechanisms of resistance to suicide switch, we performed a genome-wide CRISPR/Cas9 gene knockout screen (Fig. 3A). Jurkat T cells were sequentially transduced to express Cas9, ICAD-CAD suicide switch, and a sgRNA library^38^. Comparing lenalidomide-treated cells to untreated cells, the perturbation of two genes, *DFFB*, which encodes for the CAD protein, and *CRBN* conferred relative resistance (Fig. 3B). As lenalidomide is a molecular glue that recruits neosubstrates to the CRL4_CRBN_ E3 ubiquitin ligase^39,40^, the requirement of *CRBN* for lenalidomide-induced protein degradation was expected. To evaluate the effect of *CRBN* perturbation on suicide switch function, CRBN depletion with two sgRNAs was confirmed by Western blot. *CRBN* targeting conferred relative resistance to ICAD-CAD suicide switch as compared to a control sgRNA (Fig. 3C-D). Notably, neither perturbations in DNA damage response nor cell death pathways emerged as putative escape mechanisms in these experiments. In summary, by using a direct effector of genome cleavage as a suicide switch effector, we only detected anticipated genetic mechanisms of escape from the ICAD-CAD suicide switch that reflect a mechanism based on CAD overexpression and lenalidomide-induced destabilization of ICAD.

**Figure 3.**
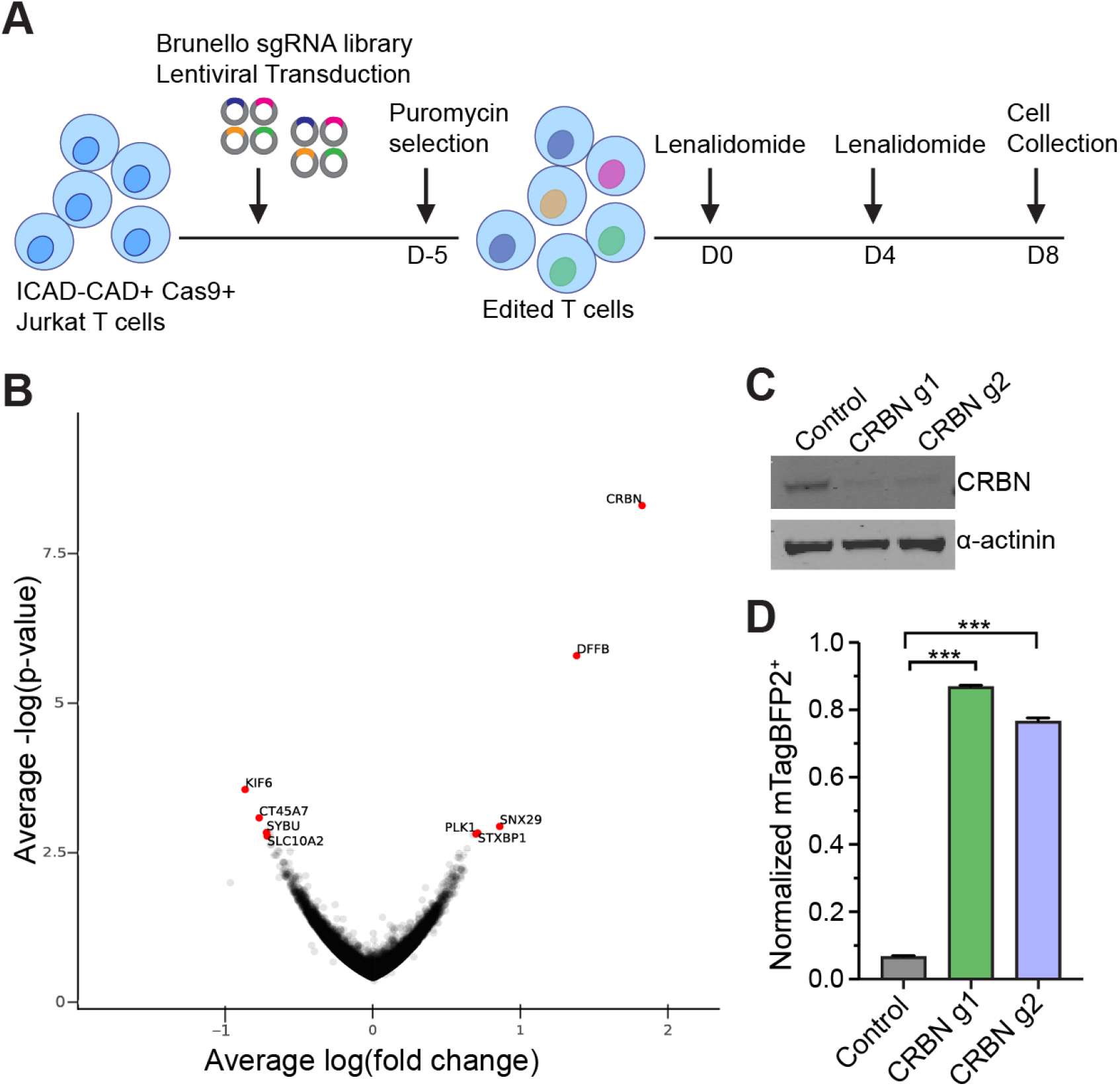
Genome-wide analysis of ICAD-CAD suicide switch resistance mechanisms (A) Schematic of the genome-wide CRISPR screen using Brunello sgRNA library. (B) Log(fold change) in gene expression summarizing CRISPR/Cas9 knockout guide abundance in ICAD-CAD+ Cas9+ Jurkat T cells after lenalidomide treatment. (C) Western blot of *CRBN*-targeted gRNA expressing Jurkat cells. Non-targeting gRNA was used as a control. (D) Normalized mTagBFP2 intensity of ICAD-CAD+ Cas9+ Jurkat T cells transduced to express control or *CRBN-*targeted gRNAs with 1 µM lenalidomide treatment (technical triplicate, mean ± SD, two-tailed student’s t-test. \**P* < 0.05, \*\**P* < 0.01, \*\*\**P* < 0.001)

### ICAD-CAD CAR T cells

CAR T cell therapies are associated with severe T cell hyperactivation syndromes that in rare cases can be refractory to existing management approaches. We hypothesized that dual vector co-delivery could enable modular CAR and ICAD-CAD suicide switch function. To this end, we co-transduced primary human T cells to express a CD19-targeting CAR and the ICAD-CAD suicide switch or vector control. Short-term lenalidomide treatment resulted in ICAD-CAD+ cell depletion (Fig. 4A-B). In a multi-day live-cell imaging co-culture assay where we assessed for high (1:5) and low (1:1) tumor burdenFACS-purified control CAR19 and ICAD-CAD CAR19 T cells exhibited comparable anti-tumor activity and CAR T cell proliferation. With lenalidomide, ICAD-CAD CAR19 T cell growth and effector functions were not detected (Fig. 4C and S5A). Their cytokine profiles showed comparable IFNɣ production with CAR19 and ICAD-CAD CAR19 T cells, while IL2 production in ICAD-CAD CAR19 T cell varied between donors (Fig. 4D and S5B). As expected, lenalidomide treated and CAR-stimulated T cells had significantly increased INFɣ levels, a characteristic of immunomodulatory imide drugs^41^. In summary, when CAR19 and ICAD-CAD were co-delivered on separate lentivectors, we observed robust *in vitro* activity from the CAR and the suicide switch.

**Figure 4.**
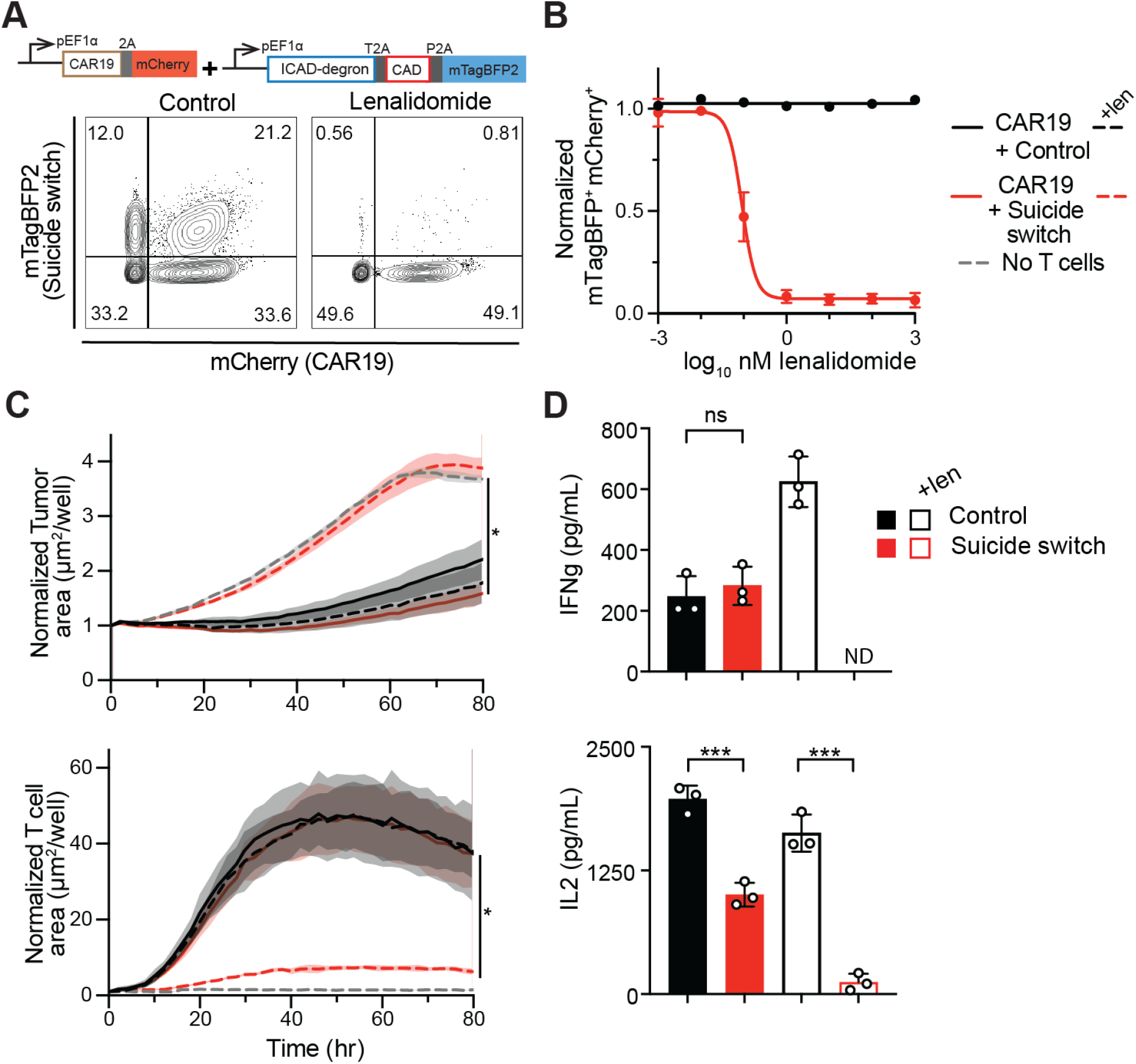
CAR and ICAD-CAD function with dual vector delivery (A) Vector map and abundance of ICAD-CAD suicide switch and anti-CD19 CAR co-transduced cells after short term treatment with 1 µM lenalidomide treatment. (B) Normalized mCherry/mTagBFP2 fluorescence ratio in primary human T cells after 20 hr treatment with varying lenalidomide concentrations. (C) Normalized tumor area and T cell area from multi-day live-cell imaging co-culture studies. NALM6 cells were co-cultured in a 1:5 ratio with co-transduced CAR19-suicide switch and CAR19-control primary human T cells. CAR19-suicide switch and CAR19-control cells were treated with 1 µM lenalidomide. Normalization was performed with the first time point. (D) Soluble cytokine bead array measurements performed with incucyte supernatants. Panel B performed with 2HD, mean ± SD. Panels C-D with 1HD, technical triplicate, mean ± SD. Panel C analyzed with 2-way ANOVA with Šídák’s multiple comparisons test at endpoint and panel D with ordinary one-way ANOVA with Tukey’s multiple comparisons test. \**P* < 0.05, \*\**P* < 0.01, \*\*\**P* < 0.001). Donor 3 data shown in Figure S4.

## DISCUSSION AND CONCLUSION

Here we present a fully human sequence-based suicide switch based on the overexpression of human ICAD and CAD genes, controlled by the clinically approved and broadly available drug lenalidomide. Mutations in the CAD CIDE domain enabled long-term expression in primary human T cells. Once triggered, the ICAD-CAD suicide switch induced more complete cell depletion than iCasp9. Mechanistically, the ICAD-CAD switch was compatible with both molecular glue and PROTAC degron systems, and cell death was correlated with CAD protein abundance. From whole-genome CRISPR screens, the only genetic perturbations associated with escape were on mechanism. ICAD-CAD and a CAR were co-delivered in a single lentivirus, but the modular function of each element was not preserved, consistent with general challenges in large and multi-transgene co-delivery^42^. While dual-lentivector co-delivery required co-selection by cell sorting, this approach enabled high performance from the CAR and suicide switch.

Existing suicide switch designs have strengths and weaknesses. The iCasp9 is clinically validated in trials of allogeneic T cell or CAR T cell therapy^12^. However, iCasp9 relies on AP1903, a chemical dimerizer not yet approved for clinical use, as well as a relatively narrow expression range with acceptably low drug-independent switch leakiness and also sufficient drug-dependent cell elimination^13^. Similar rapamycin-based systems may overcome at least the first of these challenges^43^. The Herpes Simplex Virus thymidine kinase transgene induces cell death upon addition of the prodrug ganciclovir and conversion into an active DNA replication inhibitor, and is also clinically validated as a strategy to control GvHD after donor lymphocyte infusion^44^. However, T cell responses directed at the viral transgene were frequently detected, and this immunogenicity limited in vivo persistence^9^. The expression of surface epitopes for monoclonal antibody therapeutics such as rituximab^10^ or cetuximab^11^ rely on cell-extrinsic effector mechanisms that may be diminished in people with advanced cancers, and also acts by exposing patients to the side effect profile of the paired antibody therapeutic. The more recently described metabolic *UMPS*-knockout auxotrophy suicide switch is uniquely transgene-free^45^, although it does require non-trivial additional manufacturing cost and complexity of genome editing. Each of these systems is likely to align with different translational use cases.

The best functional attributes of the ICAD-CAD suicide switch were 1) its fully human sequence-based design, 2) use of an approved, off-patent controller drug, and 3) best-in-class inducible cell depletion without detectable leakiness of CAD activity. Favorable performance characteristics compatible with high lentiviral expression were likely enabled by the chaperone/inhibitor mechanism inspired by native cell death machinery, whereby ICAD cages newly translated CAD proteins in inert heterodimers^17,18^. Once triggered with lenalidomide, this led to a sufficient concentration of uninhibited CAD protein to kill >95% of transduced cells. The principal limitation of the ICAD-CAD switch was its 2.2 kilobase transgene size, which is larger than other safety systems and currently difficult to co-deliver with effector elements in a single viral vector. Therefore, with respect to applications, ICAD-CAD may be further enabled by emerging delivery modalities with larger transgene carrying capacity^46^ or sort-free dual-selection technologies^47^. ICAD-CAD may already be well-positioned for diverse iPSC-based applications that are compatible with stepwise genetic engineering and selection of a master cell product. Alternative pairs of chaperone/inhibitor and cell death effector might also be mined or engineered for suicide switch development to exploit the degradable cage design.

Beyond engineered T cells, the CAD and ICAD stoichiometric pair as well as the requisite CRL4_CRBN_ E3 ligase components are almost ubiquitously expressed across cell types^48^. Therefore, while re-optimization may be necessary for deployment to additional cell types, the ICAD-CAD suicide switch has broad potential to safeguard diverse gene- and cell-based therapies, or act as a remote controlled warhead for diseased cell state-selective synthetic sensors^49,50^.

## MATERIALS AND METHODS

### CAD-ICAD sequences

CAD variants were synthesized and cloned into lentiviral vectors. Each vector contained an ICAD-degron, CAD variant, and mTagBFP2 fluorescent reporter. CAD sequence from O76075-1 (Uniprot). ICAD sequence from O00273-1 (Uniprot). ICAD-degron fusion composed of ICAD, GSGSGSGSGG linker, and superdegron^28^. CAR19 sequence as previously published^51^.

### Cell lines

The cell lines used in this study were HEK293 (female, fetal), Jurkat (male, 14 year-old), SupT1 (male, 8 year-old), NALM6 (male, 19 year-old), and K562 (female, 53 year-old) all obtained from ATCC. Cells were maintained at 37°C with 5% carbon dioxide and cultured in R10 medium containing RPMI 1640 medium with GlutaMAX and HEPES supplemented with 10% fetal bovine serum (FBS), penicillin, and streptomycin unless otherwise indicated. Mycoplasma-free cultures were confirmed routinely by PCR.

### Lentivector cloning

Entry vectors were linearized by restriction enzymes (New England Biola). Double stranded DNA fragments synthesized and were introduced into the vectors by T4 ligation (Thermo Fisher Scientific EL0011) according to manufacturers’ instructions. The products were transformed into STBL3 bacteria (Thermo Scientific, C737303). Sequences were confirmed by Sanger or nanopore sequencing.

### Lentiviral production

HEK293 cells cultured in r10 medium were used to produce lentiviral particles. For each mL of HEK 293 cultures, 34 mL Optimem, 0.27 mg transfer plasmid, 0.45 mg VSV-G (Addgene plasmid # 14888, produced by Dr. Tannishtha Reya), 0.18 mg pSPAX2 (Addgene plasmid # 12260, produced by Dr. Didier Trono) and 3 mL of Fugene HD were used. For cell lines, the supernatant containing lentiviral particles was collected 48 hours post transfection, filtered using a 0.45 µmM PVDF membrane and either added to target cells or stored at -80°C. For application in primary human T cells, the supernatant post 48hr of transfection was concentrated by ultracentrifugation at 25,000 RPM in a SW28 rotor, resuspended in approximately 1/100 of the starting volume in PBS, and cryopreserved at -80 C.

### Primary human CAR T cell transduction and culture

Human T cells were purified as mentioned previously^51^. Cells from at least 2 healthy donors were used in primary T cell experiments. In dual vector CAR-suicide switch T cell studies, cells from 3 donors were used.

Primary human T cells were thawed in R10 medium with 20 IU/mL recombinant human IL-2. They were activated on day 0 post thaw using CD3/CD28 Dynabeads at a 1:3 cell:bead ratio with a starting cell density of 1e6/mL. Cells were transduced on day 1 with lentivirus at a multiplicity of infection of 5. In dual lentiviral transductions, the second virus was added on day 2. From day 3, the R10 media containing recombinant-IL2 were doubled every alternate day. CD3/CD28 beads were removed from the cells using a magnet on day 7. Between day 12-14, lentiviral transduction efficiency was determined by flow cytometry. The cells were either applied to functional assays or cryopreserved in 90% FBS and 10% DMSO.

### ICAD degradation assay

Green fluorescent protein (GFP) was fused to degron-tagged ICAD to form a GFP-ICAD-degron fusion protein. GFP-ICAD-degron DNA was synthesized and cloned into lentiviral vectors. Jurkat T cells were transduced to express GFP-ICAD-degron construct. Flow cytometry (Fortessa X-20, BD Biosciences) was used to assess average GFP fluorescence intensity and ICAD abundance prior to lenalidomide treatment. ICAD degradation was assessed by the decrease in GFP reporter intensity following lenalidomide treatment by flow cytometry.

### Immunoblot analysis

Cell lysates were collected using ice-cold RIPA lysis buffer (Thermo Scientific, 89900) and freshly added protease-phosphatase inhibitor cocktail (Thermo Scientific, 78440) following the manufacturer’s protocol. Next, 30 µg of proteins were separated with 10% Bis-Tris gels (Invitrogen, NW00105BOX) then transferred onto nitrocellulose membranes (Thermo Scientific, 88018) as per the manufacturer’s instruction. After the transfer, membranes were blocked by a blocking buffer (LICORbio, 927-70001) and then stained using primary antibodies (1:1000-1:5000 dilution) followed by secondary antibodies (1:3000 dilutions). The blots were developed using a near-infrared (NIR) fluorescence imaging system (Odyssey CLx, LICORbio).

Primary antibodies used in the study includes ICAD (R&D Systems, AF4679), CAD (R&D Systems, MAB6289), FKBP12 (Abcam, ab58072), CRBN (Novus Biologicals, NBP1-91810), α-actinin (Cell Signaling Technology, 6487S) and cyclophilin B (Cell Signaling Technology, 43603S). Fluorescent labeled secondary antibodies were used (LICORbio, 926-68074, 926-32211, 926-68072 and 926-32214). All the immunoblots were carried out in duplicates.

### Switch functionality assay

Fluorescent mTagBFP2 was included in the ICAD-CAD suicide switch constructs as a reporter of suicide switch expression. Depletion of mTagBFP2 fluorescence was indicative of suicide switch-mediated apoptosis. Cells were stained with Annexin V (BioLegend, 640908, 422201), γ-H2AX (BD Biosciences, 560445) or 7-AAD (Thermo Fisher Scientific, 00-6993-50) to assess DNA damage and cell death respectively. Flow cytometry (Fortessa X-20, BD Biosciences, Aurora, Cytek Biosciences) was used to quantify the abundance of γ-H2AX (intracellular flow), 7-AAD, and mTagBFP2 reporter. iCasp9 suicide switch expression was measured by anti-NGFR expression (BioLegend, 345131). For multicistronic suicide switch-CAR19 cells, cells were stained with myc tag antibody (Cell Signaling Technology, 2279S) or recombinant human CD19 Fc Chimera Protein (R&D Systems, AFG9269-020) and quantified by flow cytometry to assess CAR19 abundance. Primary human T cells expressing dual vector suicide switch and CAR19 were purified by FACS (Sony SH800, Sony MA900) and the dual positive mCherry, mTagBFP2 intensity was estimated by flow cytometer.

### Proliferation assays

A Carboxyfluorescein succinimidyl ester (CSFE) assay was performed on Jurkat cells transduced with lentiviral vectors containing the different CAD variants. Cells were stimulated with aAPC K562 target cells in 1:3, 1:1, or 3:1 ratios or in a 1:1 ratio with CD3/28 beads. Cells were expanded in either R10 or R10 with 1 µM lenalidomide over a period of 5 days. CAD functionality was assessed by flow cytometry analysis of normalized abundance of expression mTagBFP2 reporter.

Primary human T cells expressing ICAD-CAD, iCasp9 suicide switches and empty vector mTagBFP2 expressing control were normalized and co-cultured in a 1:1 ratio with target irradiated aAPC+ K562 cells^52^. Primary human T cells expressing Suicide switch-CAR19 were co-cultured in a 1:1 ratio with target irradiated CD19+ K562 cells. Cells were expanded in R10 media or R10 media with 1 µM lenalidomide. Cells were re-stimulated on a weekly basis in a 1:1 ratio of cells to irradiated k562 cells. Media was doubled at day 3 following stimulation with K562 cells. Flow cytometry of fluorescence proteins was used to assess engineered cell abundance.

### Genome-wide CRISPR Screen

To perform the genome-wide CRISPR screen for modifiers of the suicide-switch sensitivity, we transduced Jurkat cells expressing the CAD-iCAD suicide switch-BFP with lentivirus containing Cas9 and the Brunello sgRNA library^38^ and performed selection with puromycin for 5 days. The suicide switch was activated with 1uM lenalidomide and the cells were grown for an additional 8 days. gDNA was isolated and integrated sgRNA sequences were amplified by PCR and sequenced on an Illumina Miseq using MiSeq Reagent Kit v2 (50 cycles, MS-102-2001)^53^. Enrichment or depletion of sgRNAs targeting a specific gene were calculated using the hypergeometric distribution tool in the CRISPR Gene Scoring Tool (https://portals.broadinstitute.org/gpp/public/analysis-tools/crispr-gene-scoring-help) from the Broad Institute.

### Functional CAR T cell assays

For live cell analysis of CAR T and tumor cells co-culture, tissue culture plates were coated with CD71 antibody (BioLegend, 334102) diluted in PBS the day before the experiment. The next day, primary human T cells expressing CAR19, multicistronic suicide-CAR19 or dual vector suicide switch were normalized and co-cultured at defined cell ratios with 1e5/mL NALM6 tumor cells. Cells were loaded on the Incucyte ZOOM instrument for real-time fluorescent quantification of cells. NALM6 tumor area was assessed by GFP fluorescence. CAR abundance was assessed by mCherry fluorescence. 1 µM lenalidomide was administered at the time of setup to induce suicide switch-mediated apoptosis and ablate CAR activity.

To estimate cytokine production, supernatants from the above co-cultured experiments were collected at the endpoint. The experiment was carried out using a cytometric Bead Array as per the manufacturer’s protocol (BD Biosciences, 561666). The fluorescence based read out was measured using a flow cytometer (Fortessa X-20, BD Biosciences)

### Quantification and statistical analysis

All graphs were plotted and statistically analysed in GraphPad Prism 11. Unless otherwise stated, data are represented as mean ± standard deviation (SD). Statistical tests performed are indicated in figure legends. Significance for a *P*-value was considered <0.05. 3D structures were visualized in ChimeraX. Adobe Illustrator 2025 was used to generate the figures.

## ACKNOWLEDGEMENTS

We thank members of the Jan and Maus laboratories for feedback and discussion. The work was supported by a Krantz Family Center for Cancer Research Award to M.V.M., R.M., and M.J..

## AUTHOR CONTRIBUTIONS

D.S. and M.J. conceptualized and led the studies. D.S., W.L. and M.J. developed and validated the suicide switch. D.S., W.L., T.K., M.V.M., and M.J. designed and performed the comparison to iCasp9. D.S., V.K., A.S., M.S. and M.J. designed and validated the dual degron ICAD-CAD experiments. N.K, A.C. and R.M. designed and performed the whole genome CRISPR screen and analysis while D.S., W.L., and M.J. validated the top hits. D.S., W.L., J.K., I.C.L. and M.J. designed and performed CAR T cell experiments. D.S., W.L., J.E., J.K., S.M.C., E.S., G.K. and M.J. performed the vector cloning experiments. D.S., W.L., and M.J. wrote the manuscripts with inputs from all the authors. M.J. supervised and provided funding for this work.

## ASSOCIATED CONTENT

Supplementary information contains four figures.

