## Supplemental Figure 1-5 for "A Lenalidomide-Inducible Suicide Switch for Gene- and Cell-Based Therapy"

### SUPPLEMENTARY INFORMATION

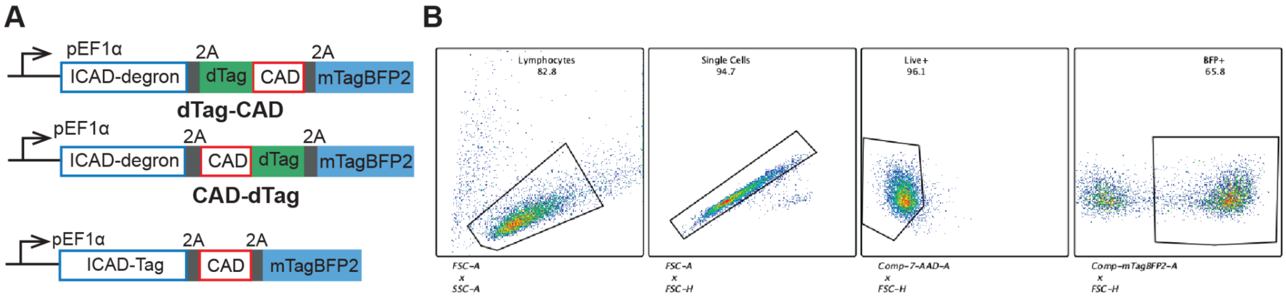


**Figure S1:** Construct strategy and Flow gating. (A) Vector map of dTag-CAD, CAD-dTag and ICAD-dTag-CAD suicide switches. (B) Representative Flow gating for BFP+ suicide switch cells.

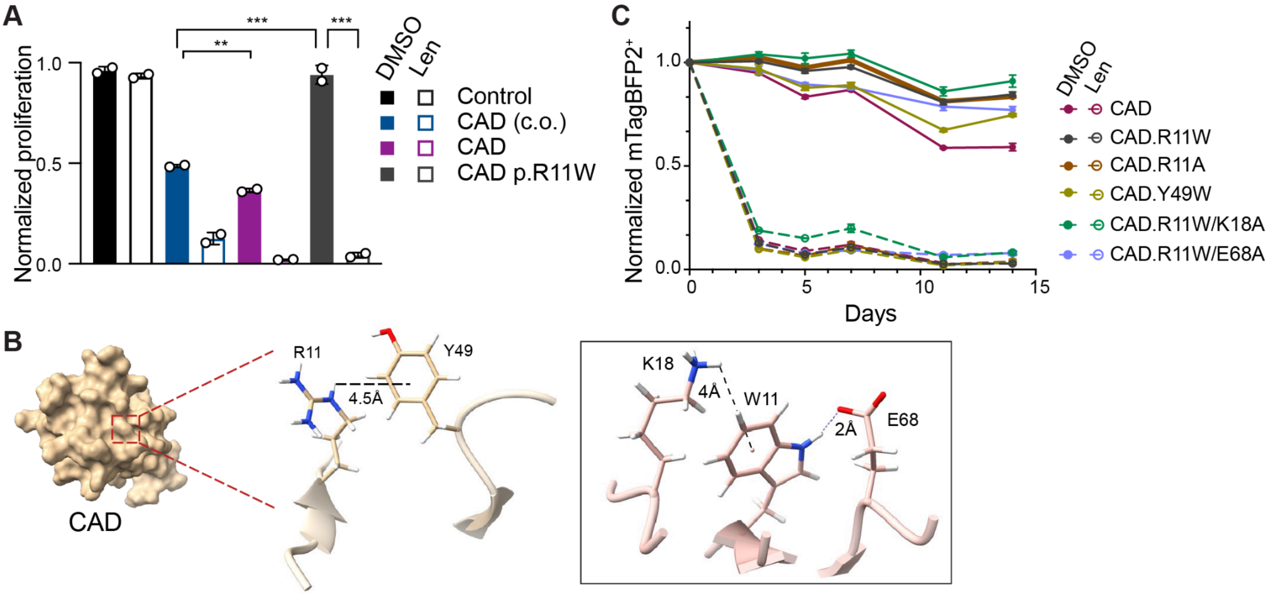


**Figure S2:** ICAD-CAD suicide switch with variation in CAD transgene sequence
(A) Carboxyfluorescein succinimidyl ester (CFSE) assay of different CAD transgenes including codon optimized (c.o.) CAD in primary human T cells with and without 1 μM lenalidomide treatment (1HD, technical duplicate, mean ± SD, ordinary one-way ANOVA with Tukey's multiple comparisons test. *$P$ < 0.05, **$P$ < 0.01, ***$P$ < 0.001). (B) NMR structure of CAD CIDE domain (PDB 1IBX) depicting interactions with R11 side chain. The box to the right represents the same with the side chain of W11. R→W mutation at position 11 was performed *in silico*. (C) 14 day proliferation assay of CAD-variant suicide switches expressed in Jurkat T cells. Cells were treated with 1 μM lenalidomide (technical triplicate, mean ± SD).

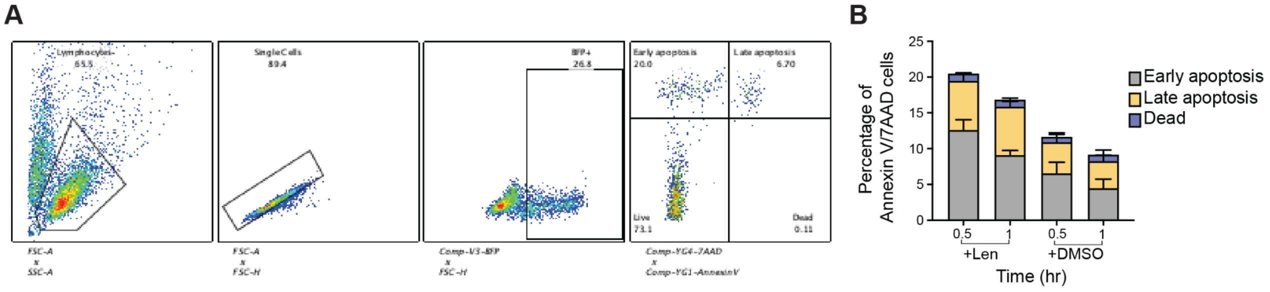


**Figure S3:** Annexin V/7AAD staining (A) Representative Flow gating. (B) Percentage of Annexin V/7AAD in primary human T cells expressing ICAD-CAD suicide switch treated with 1 μM lenalidomide and at indicated time points.

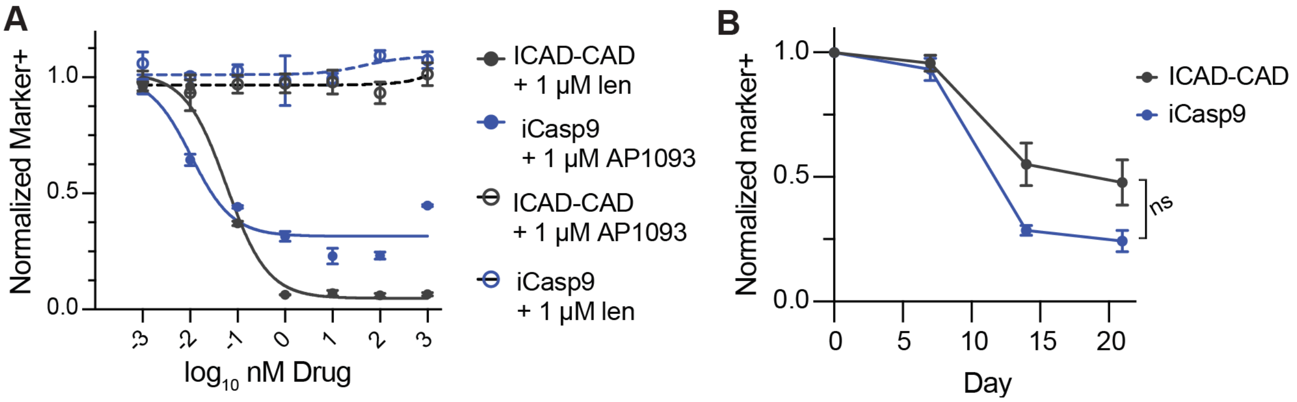


**Figure S4:** Comparison of ICAD-CAD with iCasp9 suicide switch
(A) Normalized marker+ plot of ICAD-CAD (mTagBFP2+) and iCasp9 (NGFR+) in primary human T cells after treatment with varying lenalidomide or AP1903 concentrations (1HD, technical duplicate, mean ± SD). (B) Long-term proliferation assay of primary human T cells transduced to with the suicide switches (1HD, technical triplicate, mean ± SD, 2-way ANOVA with Šídák's multiple comparisons test at day 21, *$P < 0.05$, **$P < 0.01$, ***$P < 0.001$).

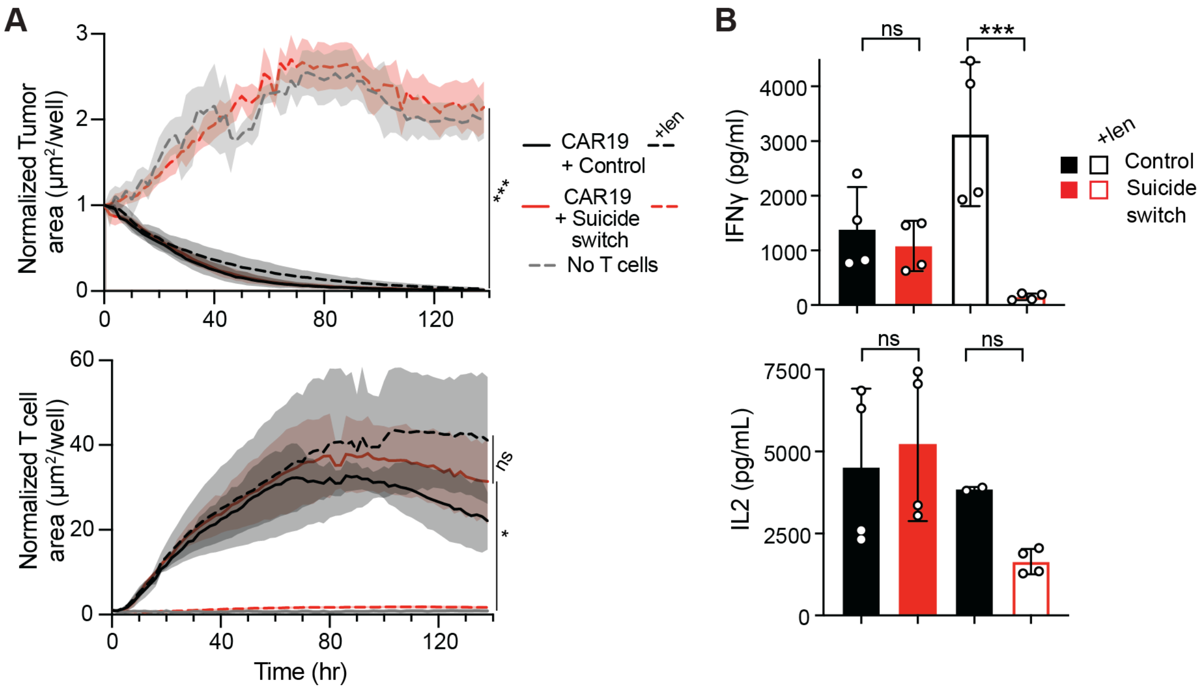


**Figure S5:** CAR and ICAD-CAD function with dual vector delivery

(A) Normalized tumor area and T cell area from incucyte studies. NALM6 cells co-cultured in a 1:1 ratio with co-transduced CAR19-suicide switch and CAR19-control primary human T cells. Cells were treated with 1 μM lenalidomide. (C) Soluble cytokine bead array measurements performed with the incucyte supernatant. All experiments carried out with 2HD, technical duplicate, mean ± SD. Panel A, analyzed with 2-way ANOVA with Šídák's multiple comparisons test at endpoint and panel B with ordinary one-way ANOVA with Tukey's multiple comparisons test. *$P < 0.05$, **$P < 0.01$, ***$P < 0.001$).
